# Does alpha phase modulate visual target detection? Three experiments with tACS phase-based stimulus presentation

**DOI:** 10.1101/675264

**Authors:** Tom A. de Graaf, Alix Thomson, Shanice E.W. Janssens, Sander van Bree, Sanne ten Oever, Alexander T. Sack

**Author notes:** Corresponding author contact details, Oxfordlaan 55, 6229 EV, Maastricht, the Netherlands, Section Brain Stimulation and Cognition, Department of Cognitive Neuroscience, Faculty of Psychology and Neuroscience, Maastricht University, Maastricht, the Netherlands.

## Abstract

In recent years the influence of alpha (7-13 Hz) phase on visual processing has received a lot of attention. Magneto-/encephalography (M/EEG) studies showed that alpha phase indexes visual excitability and task performance. If occipital alpha phase is functionally relevant, the phase of occipital alpha-frequency transcranial alternating current stimulation (tACS) could modulate visual processing. Visual stimuli presented at different pre-determined, experimentally controlled, phases of the entraining tACS signal should then result in an oscillatory pattern of visual performance. We studied this in a series of experiments. In experiment one, we applied 10 Hz tACS to right occipital cortex (O2) and used independent psychophysical staircases to obtain contrast thresholds for detection of visual gratings in left or right hemifield, in six equidistant tACS phase conditions. In experiments two and three, tACS was at EEG-based individual peak alpha frequency. In experiment two, we measured detection rates for gratings with (pseudo-)fixed contrast levels. In experiment three, participants detected brief luminance changes in a custom-built LED device, at eight equidistant alpha phases. In none of the experiments did the primary outcome measure over phase conditions consistently reflect a one-cycle sinusoid as predicted. However, post-hoc analyses of reaction times (RT) suggested that tACS alpha phase did modulate RT in both experiments 1 and 2 (not measured in experiment 3). This observation is in line with the idea that alpha phase causally gates visual inputs through cortical excitability modulation.

## Introduction

The visual hierarchy is one of the most extensively investigated systems in the human brain, but its communication mechanisms remain unclear. In recent years, studies have increasingly focused on the role of oscillatory mechanisms in successful visual processing (e.g. Gallotto, Sack, Schuhmann, & de Graaf, 2017). Oscillations can be described in terms of frequency, amplitude, and phase. In humans, magneto- or electroencephalography (M/EEG) can be used to measure such oscillatory activity, arising from synchronized neuronal ensembles (Berger, 1929).

Neuronal activity in parieto-occipital cortex oscillating at alpha frequency (7-13 Hz) seems particularly important for vision and visual attention (e.g. Klimesch, Sauseng, & Hanslmayr, 2007; Mathewson et al., 2011). The power of posterior alpha activity decreases when opening the eyes, allowing processing of visual inputs (Berger, 1929; de Graaf, Duecker, Stankevich, ten Oever, & Sack, 2017). Alpha power decreases in occipito-parietal cortex contralateral to the attended hemifield (Romei et al., 2008a; Sauseng et al., 2005; Thut, Nietzel, Brandt, & Pascual-Leone, 2006). Furthermore, alpha power relates to visual excitability: phosphenes induced by occipital pulses of transcranial magnetic stimulation (TMS) require lower stimulation intensity in participants with lower levels of resting state alpha power (Romei, Rihs, Brodbeck, & Thut, 2008b), and on trials with lower alpha activity right before TMS pulses (Romei et al., 2008a). Visual stimuli also are more readily detected on trials with lower posterior alpha power (Lange, Oostenveld, & Fries, 2013; van Dijk, Schoffelen, Oostenveld, & Jensen, 2008).

As is the case for alpha power, there is concrete evidence that the phase of ongoing alpha oscillations affects visual processing (Callaway & Yeager, 1960; Dustman & Beck, 1965). Hits (i.e. detection) or misses of flashes of light at luminance threshold were associated with different pre-stimulus alpha-theta phase distributions (Busch, Dubois, & VanRullen, 2009). Mathewson et al. (2009) showed in a metacontrast masking paradigm that posterior alpha phase could predict both visual task performance and EEG activity elicited by the visual stimulus. TMS-induced phosphene perception also depends on alpha phase, suggesting that alpha phase indexes visual excitability (Dugué, Marque, & VanRullen, 2011). TMS pulses prior to visual stimuli can abolish their perception, possibly through effects on alpha oscillations (de Graaf, Duecker, Fernholz, & Sack, 2015; de Graaf, Koivisto, Jacobs, & Sack, 2014; Jacobs, de Graaf, & Sack, 2014).

Correlational studies as discussed above are highly informative, but also constrained. Assigning trials to power or phase bins post-hoc, based on naturally occurring oscillations, limits which brain regions, frequencies, or outcome measures we can evaluate (ten Oever, de Graaf et al., 2016). Bringing power and/or phase of oscillations in specific brain regions under experimental control would enable additional research questions. Moreover, it has been argued that turning such oscillatory measures into independent variables provides solid ground for an evaluation of their causal role in particular functions (Herrmann, Strüber, Helfrich, & Engel, 2015). Such experimental control can be achieved through rhythmic sensory (de Graaf et al., 2013; Mathewson, Fabiani, Gratton, Beck, & Lleras, 2010; Mathewson et al., 2012), or rhythmic non-invasive brain stimulation (Thut, Schyns, & Gross, 2011a). Phase alignment and amplification of neuronal oscillations by an external oscillator has been called entrainment (Thut, Schyns, & Gross, 2011a).

Entrainment of alpha oscillations has been achieved with bursts of rhythmic TMS pulses, or sustained transcranial alternating current stimulation (tACS). Alpha TMS bursts have demonstrable effects on visual task performance (Jaegle & Ro, 2014; Romei, Gross, & Thut, 2010) and local alpha power measured by EEG (Thut et al., 2011b). Jaegle and Ro (2014) showed that visual target processing was affected by preceding parietal alpha TMS bursts in a time-specific manner, in line with a causal role of alpha phase. Helfrich et al. (2014) reported that occipital alpha tACS indeed entrained neuronal alpha oscillations, with consequences for visual processing.

This converging evidence suggests that both power and phase of posterior alpha oscillations are causally relevant for successful detection of visual targets. Specifically relevant for the study of alpha phase, we recently developed experimental methodology (ten Oever, de Graaf, et al., 2016) that not only allows full experimental control over tACS stimulation, but also sub-millisecond precise time-locked presentation of (multi-modal) sensory or magnetic stimuli to participants. Aside from other benefits, this setup allows tACS-phase based stimulus presentation, which turns oscillatory phase into an independent variable. In the current series of experiments, we took advantage of this implementation to present visual gratings (experiments 1/2) or LED luminance changes (experiment 3) at pre-determined phases of tACS administered to right occipital cortex. If tACS successfully entrained (phase-aligned, possibly amplified) posterior neuronal oscillations, tACS alpha phase should correspond to neuronal alpha phase. This allowed us, for example, to determine contrast detection thresholds for stimuli presented at each tACS phase, separately for the right and left visual hemifields, with psychophysical staircases running in parallel for different phase conditions. This is not possible with correlational post-hoc phase binning studies, and opens up a wide range of new studies and more precise questions to ask. Yet, looking ahead, we failed to find consistent evidence for a causal role of alpha phase in our experiments, across several attempts, in our primary outcome measures. Post-hoc analyses did support alpha phase effects on reaction times.

## Materials and Methods

We performed three related experiments (exp1, exp2, exp3), for which we indicate under each header how they differ in procedure and parameters.

### Participants

This series of experiments included 39 measurements in total. Fourteen participants including two authors (T.G., S.O.) were tested in exp1, but two were excluded prior to main analyses due to problems with hardware or task performance. Ten participants including one author (S.B.) were tested in exp2 (one participant participated in both exp1/exp2). Fifteen participants were included in exp3. All participants had normal or corrected-to-normal vision, were screened for tACS safety, and provided written informed consent. The experimental procedures were approved by the local ethics committee.

### Transcranial alternating current stimulation and controlling equipment

In all experiments, tACS was applied using a small (3 x 3 cm) electrode applied over right occipital cortex (O2 position in the international 10-20 coordinate system) and a large (5 x 7 cm) reference electrode applied over vertex (Cz position) with anterior-posterior orientation for the longer side.

In experiment 1, peak-to-peak amplitude of stimulation was determined per participant. Before the main experiment, we stimulated participants briefly with the montage, with increasing stimulation intensity, asking them each time to indicate what they found comfortable and whether they perceived phosphenes to such an extent that they might interfere with the processing of visual stimuli. This resulted in a range of stimulation intensities from 0.8 to 2 mA peak-to-peak, with twelve out of fourteen participants stimulated at 1.2 mA or higher and an overall mean intensity of 1.5 mA. At the occipital electrode, this constitutes a mean current density of 1.67 A/m2 under the electrode, which ought to be sufficient to modulate neuronal activity at that location (Herrmann 2013, Neuling 2012).

TACS was ramped up and down over 10 seconds. Stimulation duration was approximately 19 minutes for experiments 1/2, 20 minutes for experiment 3. tACS frequency was 10 Hertz (Hz) for exp1, and at individual peak alpha frequency for exp2 and exp3 (see below for determination). Also in exp 2/3, tACS was briefly applied prior to the experiment, to test tolerance for somatosensory experience. We then also checked with participants that any potential phosphenes (visual experiences caused by tACS) did not occur in task-relevant parts of the visual field. In these experiments, peak-to-peak amplitude was set to 1.5 mA by default, though if stimulation was deemed too uncomfortable or phosphenes overlapped with target locations, intensity was reduced to 1 mA peak to peak, which occurred once in exp2.

TACS was remotely controlled, using the Remote option on NeuroConn DC-STIMULATOR PLUS (neuroConn, GmbH, Ilmenau Germany). We did not correct for any minor individual DC offset, which can be introduced when using Remote tACS. Electrodes were attached using Ten20 conductive neurodiagnostic electrode gel (Weaver and Company, Aurora, Colorado, USA). We previously (ten Oever, de Graaf, et al. 2016) described our experimental setup, for which we summarize the procedures and implementation below. It involves source files created in Matlab (TheMathWorks Inc., Natick, Massachusetts, USA), loaded into custom software DataStreamer (ten Oever, de Graaf, et al., 2016). TACS signal and stimulus triggering pulses were fed through a digital-analog converter (DAC) from National Instruments (Austin, TX, USA). A standard BSC cable connected the DAC to the tACS stimulation device. The resolution of the tACS waveform in the source files was 6000Hz in experiments 1 / 2 and 4000Hz in experiment 3. Stimulus triggering pulses (see below) consisted of digital values communicated by DataStreamer to the DAC through a parallel port connection. The DAC connected to the parallel port of a stimulus PC, running PsychToolbox (Brainard, 1997; Pelli, 1997) in Matlab and reading port activity to detect incoming triggering pulses. The pulses triggered presentation of visual stimuli on a standard LCD monitor with stimulus parameters depending on pulse values, in exp1 and exp2. In exp3, pulses triggered visual stimuli not on the computer monitor, but on a custom-built LED device connected to a parallel port of the stimulus PC.

### Stimuli and tasks: experiments 1 and 2

In exp1 and exp2, visual stimuli were circular gratings presented on a grey background (∼54 cd/m2) on the gamma-corrected display. Participants were seated 57 cm from the display, with head fixated by use of a chin rest. Vertical gratings, 1.5 degrees visual angle (DVA) in diameter, presented diagonally to either lower left (left hemifield) or lower right (right hemifield) at 6.5 DVA eccentricity. Spatial frequency was 2 cycles/degree, phase was randomized each trial, edges were faded with Gaussian blur.

The 2-alternative forced-choice task of participants was always to indicate, on each trial, whether a grating had been presented in the left or right hemifield with button presses on a keyboard. Per hemifield, detection rate (proportion correct) was calculated per tACS phase condition. Since gratings were presented at peri-threshold contrast, it was important that participants were prompted to respond, also in trials where targets were missed. A black central fixation dot, otherwise presented continuously on screen, increased in brightness for 33.3 ms (‘flashed’) simultaneously with presentation of the target grating. This cue was identical across all conditions of both experiments.

In experiment 1, the contrast of visual gratings was variable across trials. In fact, separately for left and right hemifield, for each of 6 tACS phase conditions (see Figure 1 and below), we ran a *psychophysical staircase* to determine the required contrast for 80% detection rate for that hemifield/phase condition. The dependent variable across conditions was therefore the contrast threshold. The staircases used the Quest (Watson & Pelli, 1983) functionality in Psychtoolbox, a Bayesian staircase algorithm that can suggest per trial the optimal test value, and converges on final estimates based on the entire history of test values and responses in a predetermined number of test trials (40 trials per staircase in our experiment). We supplied Quest with following parameters: beta = 3.5, gamma = 0.5, delta = 0.01, tGuess = 1, tSD = 1.

**Figure 1:**
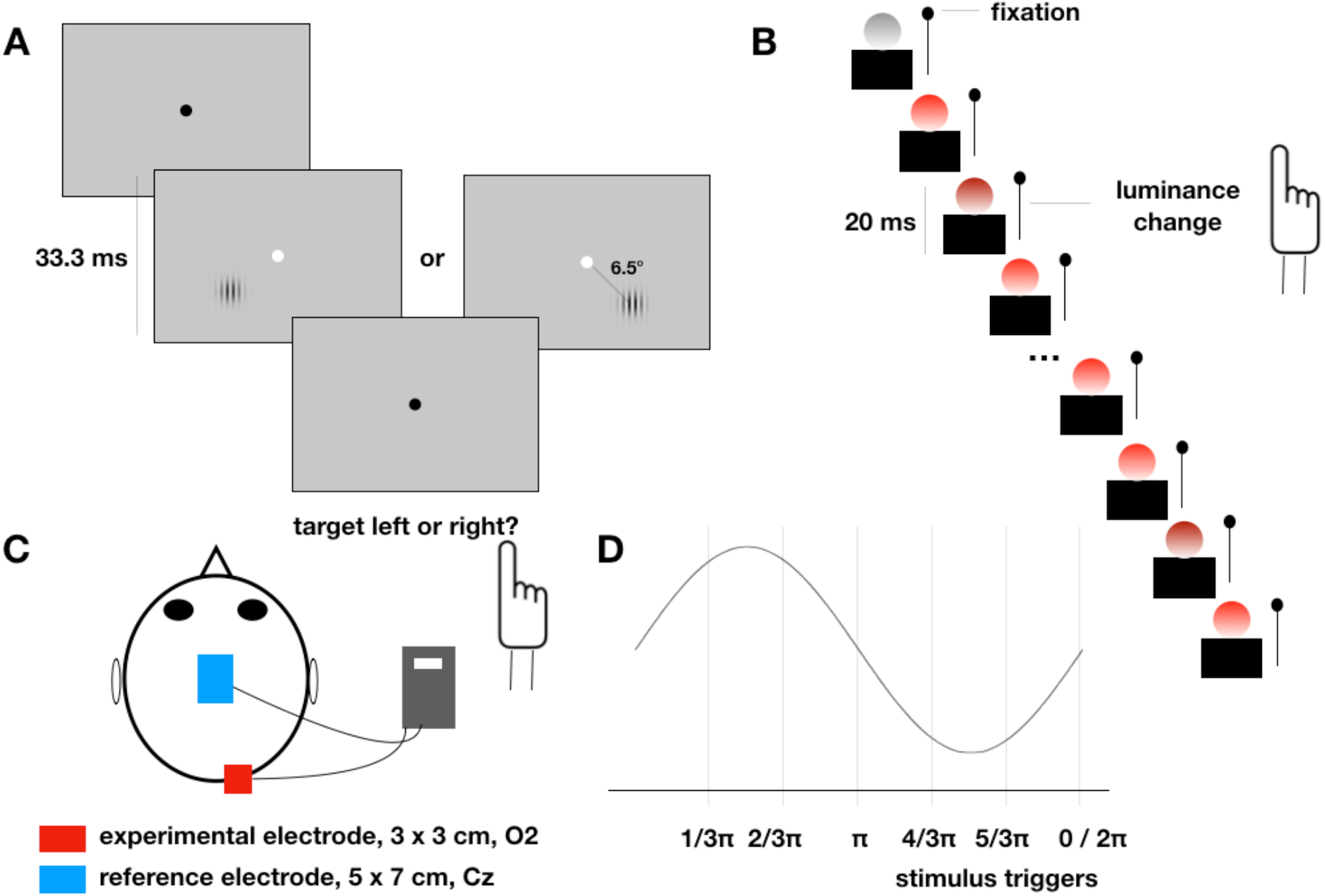
Experimental design and tasks. A. In experiments 1 and 2, participants fixated on a central black dot. Stimuli were sinusoidal gratings of calibrated contrast, presented either lower left or right of fixation per trial. Location was uncued, the fixation dot brightened to prompt a 2-alternative forced-choice response about target location.
B. In experiment 3, participants fixated a white Q-tip positioned to the upper right of an LED stimulus covered by a ping-pong ball to enlarge and diffuse the stimulus. The LED turned on to signify task start, and the LED would briefly decrease in luminance by an individually calibrated amount for 20 ms several times before turning dark again, signifying a short break. Participants responded to perceived luminance changes.
C. Participants received focal tACS to right occipital cortex (O2), with a non-focal reference electrode over vertex (Cz).
D. In all experiments, stimuli were triggered in pre-determined phases of the ongoing tACS signal. Shown are the phase conditions for experiments 1 and 2; six equidistant phases spanning one cycle. In experiment 3 there were eight equidistant phase conditions.

In experiment 2, the contrasts of visual gratings were pseudo-fixed over trials. Quest staircases, separately for left and right hemifield, determined individually calibrated contrasts to achieve 75% detection rates, prior to the main experiment. This was slightly lower than the 80% aim performance in experiment 1 to allow for performance increases over the course of the session. After all, the 80% was to be achieved by the end of sessions in experiment 1, while the 75% performance was established at the start of sessions in experiment 2. These contrasts were then fixed for the main experiment, in principle, and the dependent variable was accuracy over hemifield and tACS phase conditions. However, per hemifield, but independently of tACS phase conditions, the experiment program did keep track of task performance over time. Since we were interested in a potentially oscillating pattern of our outcome measures over tACS phase conditions, it was important that accuracy overall would not reach ceiling or floor levels. Thus, if accuracy in the most recent 5 trials reached 60% (approaching floor) or 90% (approaching ceiling), contrast was increased or decreased by 10% of its previous value, respectively.

### Stimuli and tasks: experiment 3

In experiment 3, visual stimulation was an LED, briefly changing luminance. Participants placed their heads in a chin rest in a dark environment (a dark cloth placed over their heads in a lab with lights off), and the LED device was positioned 35cm in front of them. On top of the LED device was a white stick of cotton swab (Q-tip), to serve as a fixation point to the upper right of the LED. The LED stimulus was enlarged and diffused by placing a punctured white ping-pong ball over it, to help avoid fading effects. All this resulted in a red visual stimulus with a diameter of approximately 4 cm, at a viewing distance of approximately 35 cm. From the view of the participant, the cotton swab fixation point was approximately 0.5 cm above the top right edge of the ping pong ball, placing the red target stimulus in the periphery. Participants clicked a left mouse button whenever they perceived a brief change in luminance (a ‘flicker’). The LED was in principle always on during trials, and it regularly turned off to indicate breaks. Trial events were not cued in this experiment, in contrast to experiments 1/2. This experiment was therefore a signal detection task, rather than 2-alternative forced choice task. A Quest staircase prior to the main experiment determined the decrease in LED luminance required to obtain a 45% detection rate. This luminance change was then pseudo-fixed for the main experiment. Performance on the most recent 10 trials was monitored by the experiment program, and the luminance change was increased or decreased by a factor depending on the amount of deviation from the target performance range (0.3 to 0.7 proportion of stimuli detected). The dependent variable for this experiment was thus detection rate of luminance changes, defined as the proportion of trials per tACS phase condition on which participants pressed the button within 1.4 seconds of an actual luminance change.

### tACS phase conditions and phase-locked visual presentation

Of course the core methodology across all experiments was the presentation of stimuli at predefined phases of the entraining tACS signal. This was achieved, as described previously (ten Oever, de Graaf, et al. 2016) and summarized above, by generating source files containing both the tACS stimulation and the desired timing of stimuli in relation to that tACS signal. Concretely, these source files contained values oscillating between −1 and +1, scaled by DataStreamer software to the desired tACS intensity, at a particular sampling frequency. In a secondary timeline in the source files, ‘pulses’ were coded to indicate the timing (by their position in the timeline) and parameters (by their numerical value) of visual stimuli.

In experiments 1/2, per hemifield there were 6 tACS phase conditions. This number relates to the refresh rate of our display, which is 60Hz and therefore can present six frames per 100 milliseconds (ms). Since one cycle of 10 Hz lasts 100 ms, our ‘sampling resolution’ using this display was limited to six phases of a 100 ms cycle. In the source files, there were therefore 12 possible ‘pulse values’. 1 through 6 indicated that a stimulus should be presented in the left hemifield, 7 through 12 indicated a stimulus should be presented in the right hemifield. Taking 1 through 6 as our example; each of these values was presented precisely time-locked to always the same phase of tACS, spanning one full cycle. Thus, in radians, ‘1’ always at 1/6*2pi, ‘2’ always at 2/6*2pi, etc. In experiment 1, each numerical value, directly reflecting one tACS phase condition, was associated with its own psychophysical staircase. At the end of each trial, the participant response was processed, the next contrast value to be tested was determined by Quest, and the visual grating was prepared for the next trial of that particular phase condition and hemifield.

Between trials, the experiment program was continuously scanning for incoming inputs. As soon as a triggering pulse was received, the program would as quickly as possible display the associated visual grating on screen along with the central response cue. This presentation of gratings whenever a triggering pulse was received, was inevitably not instant. With a 60Hz display, even if computational processing were instant, the time from incoming trigger to actual display would be anywhere between 0 and 16.7 ms, if the stimulus was presented at the first next available frame (screen refresh). Since the internal clocks of the PC running DataStreamer and the PC presenting visual stimuli were likely not perfectly synchronized, running an experiment like this over tens of minutes would also mean that these delays were not constant. If the first next frame was ‘missed’, the delay could be slightly longer. With a period of on average 100 ms, and a corresponding separation of tACS phase conditions of 16.7 ms, these delay variations go towards measurement noise and should be kept in mind. This is however not the case in exp3, in which we used an LED device for visual stimulation explicitly to avoid such limitations. Note that, in exp2, the use of individual peak alpha frequencies for tACS introduces an additional limitation: though triggering pulses were sent at 6 equidistant phases of the tACS signal, the display and thereby actual visual presentations were still constrained to the same frame rate of 60Hz.

### Experiment parameters and procedures

#### Experiment 1

Two visual hemifields and six phase conditions resulted in a total of 12 condition cells in a 2×6 design. Per cell, we collected 40 trials using Quest staircases. Using the QuestMean function, we extracted from the staircase performance in each condition cell a final estimate of the contrast required for 80 % task performance. Visual stimulus duration was 33.3 ms, inter-trial duration was jittered around 2 seconds, and the experiment included 5 breaks of 15 seconds.

In experimental sessions, participants first received explanations of the tasks and experimental procedures. They performed calibration measurements using Quest staircases, serving also as task practice. tACS electrodes were applied, tACS was applied at 10 Hz and participant could report on tolerance and phosphene perception. Then the tACS device was set to the Remote option, the experiment program was started, which triggered DataStreamer software to start reading out its source file, containing the tACS signal and the tACS phase-locked stimulus triggers.

#### Experiment 2

The same 2×6 design as in exp1 was implemented with 40 trials per condition cell. Visual stimulus duration was 33.3 ms, inter-trial duration was jittered around 2 seconds, and breaks were offered approximately every 3 minutes. As above, participants received explanations of tasks and procedure, were screened for tACS safety, and provided written informed consent. Participants first performed two or three calibration measurements, depending on consistency of outcome, which were Quest staircases with parameters beta = 3.5, gamma = 0.5, delta = 0.01, tGuess = 0.8, tSD = 1. Using the QuestMean function, each staircase yielded a contrast level for 75% correct detection, contrast threshold results for repeated and included staircases were averaged.

Next we determined individual peak alpha frequency (IAF). We applied single EEG electrodes to positions P4, right mastoid (reference), and Fz (ground), of the international 10-20 coordinate system. Participants closed their eyes and relaxed for 150 seconds while we recorded EEG. A fast fourier transform using the FieldTrip (Oostenveld, Fries, Maris & Schoffelen, 2011; Donders Institute for Brain, Cognition and Behaviour, Radboud University Nijmegen, The Netherlands) ft_freqanalysis.m function yielded powerspectra for 5 second epochs which were then averaged. We determined the local maximum of the resulting spectrum within the alpha-band: window 7-13 Hz. The frequency corresponding to this local maximum was taken as IAF and used to build the tACS source file. The main experiment proceeded as described for exp1. In this experiment and experiment 3, a coding error resulted in identical trial order for most participants. This means that, although the order of trials (conditions) was randomized and unpredictable for participants, it was not different between participants.

#### Experiment 3

Since the visual stimulation device here was a custom-built LED device, there was only one visual field (left) and we increased the number of tACS phase conditions to eight. Per phase condition we collected 30 trials. Trials had no clear beginning or end for participants. If the LED turned on, participants knew to pay attention to the stimulus and report any perceived luminance changes. Responses could reflect false alarms, or hits. To constrain stimulus regularity/predictability, we created inter-stimulus intervals (ITI) based on a gamma distribution: each ITI was 2 + *g* seconds, where *g* was a randomly selected value from a gamma distribution with shape parameter 1 and scale parameter 1.3. Mean inter-stimulus interval was therefore 3.3 seconds, but between any two trials the interval could be up to 15 seconds. Breaks were indicated by the LED turning off, and were given every 25 trials for 20 seconds. A longer break of 120 seconds, in which we removed the black cloth covering the participant and stimulus display, and turned on the lab lights, occurred after 129 trials. Lastly, to help prevent fading and allow participants a brief moment to relax and blink their eyes, the LED light turned off for 0.8 seconds every 5 trials.

Otherwise, procedures were similar to experiments 1/2, and individual alpha frequency (IAF) was determined as in exp2. In this experiment, three Quest staircases were run to determine the LED luminance change yielding 45% detection of the stimulus. This luminance change was used as visual stimulus for the main experiment, but adapted as dictated by performance across all conditions over the course of the experiment as mentioned above.

### Preprocessing

In experiments 1 and 2, trials on which participants pressed an incorrect key (not corresponding to either of the response options) or pressed a key too late (1.4 seconds allotted for response) were excluded (in the case of staircases in experiment 1, these trials did not contribute to the online estimation procedure). Post-hoc, trials with response times below 200 ms were also excluded. In experiment 1, staircase outcomes (based on QuestMean.m function) were recomputed after removal of these trials, by rerunning the staircases with simulated responses corresponding to actual responses from the remaining, included trials.

### Analyses and statistical tests

The outcome measures differed over experiments (exp1: contrast thresholds, exp2: accuracy, exp3: detection rate), but the analyses were largely identical. After all, whichever outcome measure we used, the hypothesis was always that this outcome measure would display a one-cycle sinusoid over tACS phase conditions. This is the consequence of our hypothesis that visual task performance should oscillate along with neuronal oscillations, which should oscillate along with tACS. The applied statistical analyses were therefore designed specifically to assess whether 1) tACS phase effects occurred, and particularly whether 2) a one-cycle sinus matched the pattern of behavioral outcomes over the sampled phase conditions.

On different levels (participant level, group level), we fitted sinuses to the 6 (exp1/exp2) or 8 (exp3) datapoints based on minimization of squared errors. These sinuses had free amplitude and phase, but fixed frequency (exactly one cycle across the datapoints). The goodness of fit would be reflected in R-squared; variance in datapoints explained by the sinusoid fit. Following previous reports (Fiebelkorn et al., 2011; ten Oever & Sack, 2015; Schilberg et al., 2018), we multiplied R-squared by the variance of the best-fitting sinusoid, to obtain *relevance values*. This hybrid measure reflects both the goodness-of-fit and the extent of modulation of performance by tACS, both of which are of interest in the current context. Note that in the Results section and figures, R-squared is often reported, as a more intuitive outcome measure, but the associated P-values are based on permutation tests of these relevance values, not R-squared values.

We used permutation tests to determine the statistical significance of obtained relevance values. For each result we wanted to statistically assess, we would build a null distribution against which to test it. This null distribution was created by repeating exactly the same processing steps on the same data, but after shuffling individual trial labels. In experiment 1, since the core outcome measure was the result of a Quest staircase procedure (using QuestMean.m function), each shuffling of trial labels was followed by a recomputation of the outcome of the Quest staircase algorithm, through simulation of the responses based on (shuffled) real responses. These steps were always taken separately per participant and in exp 1/2 per visual hemifield, so only the phase-condition labels of trials were shuffled. Two thousand iterations of shuffling and re-calculation of results (e.g. individual accuracy per condition, staircase outcome, subsequent fitting-based relevance value) led to a distribution of results. The P values we report are the proportion of permuted results (i.e. null distribution) larger than the actually obtained result. We consider outcomes falling in the last 0.05 of the null distribution to be statistically significant.

These procedures apply to all the following analyses. On the group level, we performed three different analyses. 1) we phase-aligned, Z-scored, and then averaged the individual participants’ outcome measures of phase conditions into a group result pattern. We then performed the sinusoid curve fitting analysis on this group result. Significance of the resulting fit (relevance value) was evaluated by repeating this entire analysis 2000 times on the data with shuffled trial labels (permutation test as described above). For various reasons (e.g. retino-cortical transmission time, individual visual cortical anatomy) it is not a priori expected that, even in the case of successful tACS-phase modulation of visual processing, individual results should phase-align. Therefore, we used the maxima in individual outcomes over phase conditions (i.e. the phase conditions with absolute highest task performance) to phase-shift individual results. Importantly, we left out these peak values, used for the phase-alignment, from the group analysis.

2) To allow for the possibility that tACS phase could modulate performance, just not in a sinusoidal fashion, we also tested after phase-alignment the group average of the ‘up-phase’ conditions (since phase condition 2 was the ‘peak’ after phase-alignment and therefore removed: up-phase condition was the group average of phases 1 and 3) against the group average of the ‘down-phase’ conditions (group average of phases 4, 5, and 6). (In exp 3, the analogous contrasted phases were 1, 3, 4 against 5, 6, 7, 8.) This contrast was tested in a permutation test on the up-phase minus down-phase means: trial-shuffling the phase condition labels 2000 times and always repeating the full analysis procedure up to calculation of a group up-phase average minus group down-phase average. 3) These analyses effectively constitute fixed-effects analyses. Therefore, we based a final group analysis on the results of individual curve fitting analyses and associated permutation tests. For each participant, the curve fitting procedure and individual permutation test assessed to what extent the individual pattern of performance over phase conditions statistically significantly matched a sinusoid. The resulting P-value per participant, per visual field location, per dependent variable (see below) was converted to a Z-score. The resulting vector of Z-scores was tested against 0 in a one-sample t-test. A significant deviation of the mean Z-score from 0 should indicate that, over participants, the sinusoid curves consistently explained more data than expected by chance, even if effects were too small on the individual subject level to reach significance.

Importantly, a priori we planned these analyses for the following dependent variables: contrast detection thresholds in exp 1, accuracy (proportion correct) in exp2, and hit rate in exp3. However, given the inconsistent and null results for these dependent variables (see Results), we post-hoc decided also to perform and report the same analyses for the dependent variable of reaction times, which were recorded in exp1 and exp2. On the subject and condition level, mean reaction times were estimated based only on correct trials. As will be clear below, analyses did provide some support for effects of tACS phase on reaction times. This is not unexpected, but it should be kept in mind that these analyses were secondary and post-hoc.

## Results

In experiment 1 we tested the modulation of contrast thresholds for gratings in lower left or lower right visual hemifields, when presenting these gratings at six different phases of 10Hz tACS administered to right occipital cortex (location O2). In experiment 2 we tested modulation of grating detection performance (hit rates) using pseudo-fixed thresholds, by different phases of tACS at individual peak alpha frequencies (IAF). In experiment 3 we tested modulation of LED luminance change detection by eight different phases of IAF tACS. We present the results sequentially. We additionally report the outcomes when performing the same analyses on reaction times in experiments 1 and 2 (not recorded in experiment 3).

### Experiment 1

#### Individual results

The full pattern of individual results is shown in Supplementary Figures SF1 and SF2, which present the thresholds (SF1) and mean reaction times (SF2), per hemifield, across tACS phase conditions. Best-fitting one-cycle sinusoids are superimposed, figure titles provide the explained variances, as well as the P-values based on permutation tests on the relevance values of the curve-fitting approach (see Methods). Table 1 below also provides the variances explained by fitted sinusoids (R-squared) (contrast threshold in italics as the primary analysis, reaction time as post-hoc analysis) along with P-values resulting from permutation tests on the relevance values (R-squared multiplied by amplitude of best-fitting sinusoid, see Methods). Conditions in which the relevance value was statistically significant (uncorrected) are indicated in bold.

**Table 1:**
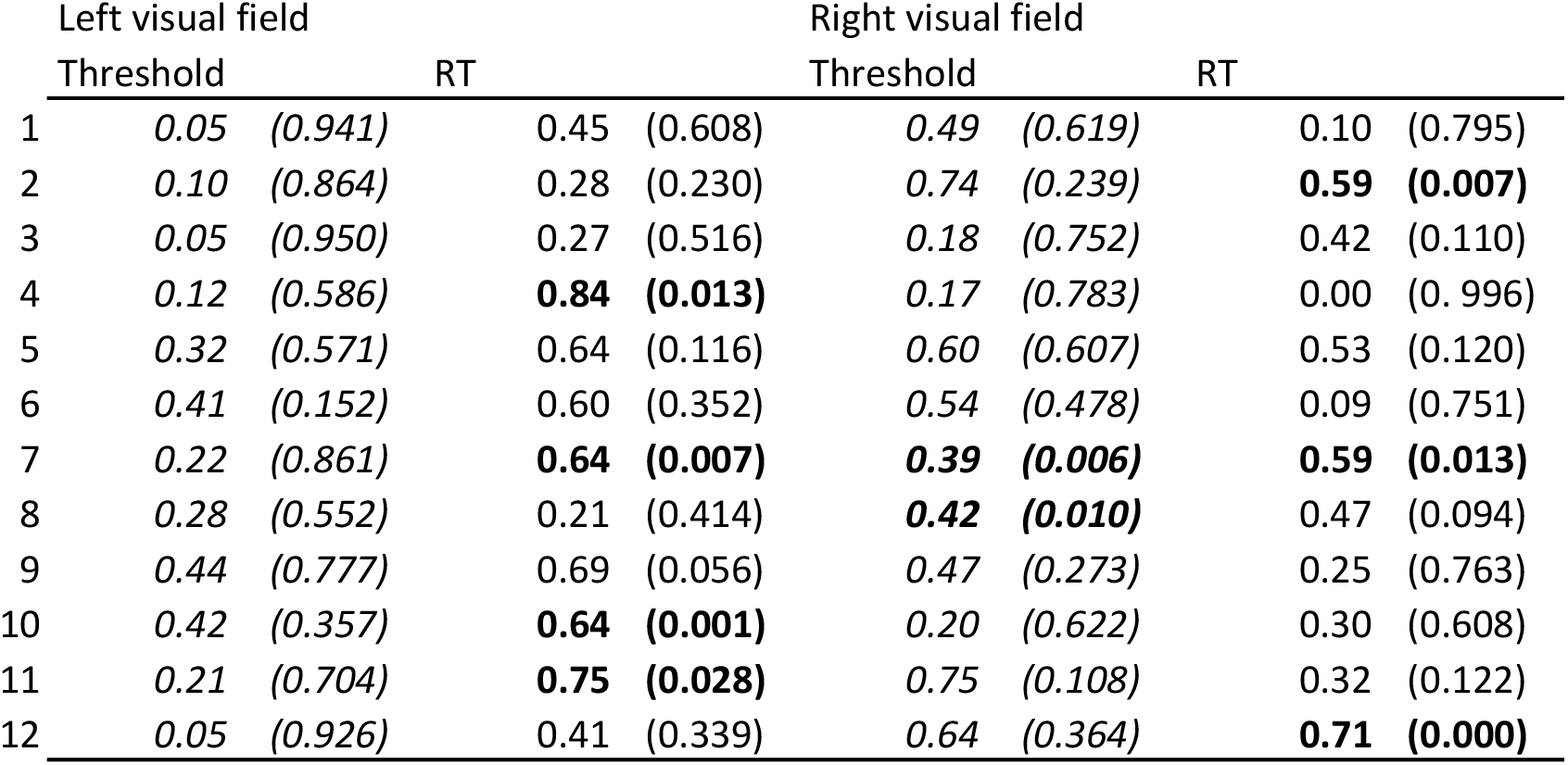
individual results. Explained variance (R-squared) with P-value resulting from permutation tests on relevance values in parentheses. Threshold analysis was the main analysis, reaction time analysis was post-hoc Bold cells are statistically significant (uncorrected)

|  | Left visual field |  |  |  | Right visual field |  |  |  |
| --- | --- | --- | --- | --- | --- | --- | --- | --- |
|  | Threshold |  | RT |  | Threshold |  | RT |  |
| 1 | 0.05 | (0.941) | 0.45 | (0.608) | 0.49 | (0.619) | 0.10 | (0.795) |
| 2 | 0.10 | (0.864) | 0.28 | (0.230) | 0.74 | (0.239) | **0.59** | **(0.007)** |
| 3 | 0.05 | (0.950) | 0.27 | (0.516) | 0.18 | (0.752) | 0.42 | (0.110) |
| 4 | 0.12 | (0.586) | **0.84** | **(0.013)** | 0.17 | (0.783) | 0.00 | (0.996) |
| 5 | 0.32 | (0.571) | 0.64 | (0.116) | 0.60 | (0.607) | 0.53 | (0.120) |
| 6 | 0.41 | (0.152) | 0.60 | (0.352) | 0.54 | (0.478) | 0.09 | (0.751) |
| 7 | 0.22 | (0.861) | **0.64** | **(0.007)** | **0.39** | **(0.006)** | **0.59** | **(0.013)** |
| 8 | 0.28 | (0.552) | 0.21 | (0.414) | **0.42** | **(0.010)** | 0.47 | (0.094) |
| 9 | 0.44 | (0.777) | 0.69 | (0.056) | 0.47 | (0.273) | 0.25 | (0.763) |
| 10 | 0.42 | (0.357) | **0.64** | **(0.001)** | 0.20 | (0.622) | 0.30 | (0.608) |
| 11 | 0.21 | (0.704) | **0.75** | **(0.028)** | 0.75 | (0.108) | 0.32 | (0.122) |
| 12 | 0.05 | (0.926) | 0.41 | (0.339) | 0.64 | (0.364) | **0.71** | **(0.000)** |

Three observations are relevant in Table 1 (and SF1/SF2). Firstly, while our main hypothesis for exp1 was that contrast thresholds should be modulated by alpha tACS phase especially in the left hemifield, not a single participant showed such modulation statistically significantly. For right hemifield targets, without multiple comparisons correction two participants showed modulation that reached significance. Secondly, sinusoids often did explain a lot of variance even if not statistically significant. This is also apparent from SF1/SF2: the goodness of fit is often visually impressive. Yet, likely due to the limited number of phase conditions and the inevitable sampling of only a single oscillatory cycle, an oscillatory pattern well matched by a sinusoid can appear by chance relatively easily. This emphasizes how important it is to think critically about the appropriate statistical procedures to test the observed goodness of fits. Thirdly, our post-hoc additional analysis evaluated modulation of reaction times (RT) by alpha tACS phase, and here specifically in left hemifield there were four participants with significant modulation (plus another approaching significance), versus three in the right hemifield.

#### Group analyses

It is difficult to credit so many individual statistical tests, or to draw generalized conclusions from them. We performed a second-level group analysis on these individual results. Per hemifield, and separately for contrast thresholds and mean RTs, the individual P-values were converted to Z-scores, which were subsequently tested against zero in a one-sided t-test across participants. Essentially this approach evaluates the extent to which the relevance values of individual participants were consistently towards the extreme (right) end of their associated permutation-based null distributions. Thus, this test might capture consistent but small effects, too weak to be significant in individual subject statistics but meaningful across the sample. This test however did not reject the null hypothesis that sinusoids fit the individual data consistently better than chance for contrast thresholds of left hemifield targets (t(11) = −2.61, P = 0.99), or right hemifield targets (t(11) = 1.44, P = 0.09). As a post-hoc analysis, it suggested that reaction times to specifically left targets might have been modulated by alpha tACS phase (left hemifield: t(11) = 3.62, P = 0.002, right hemifield: t(11) = 1.26, P = 0.12). This effect even survives Bonferroni-correction for these four tests, but keep in mind that 1) the reaction time analysis was post-hoc, and 2) the experiment was not designed with this analysis in mind. So we would like to replicate it in experiment 2, and in the other group analyses we performed on this data.

In a second group analysis, we first phase-aligned individual performance patterns, phase-shifting each pattern based on the absolute peak in performance across the six phase conditions. These peak datapoints were then excluded from further analysis. The remaining (five) phase condition results were averaged across participants, and a once-cycle sinusoid was fitted to the group average result. See Figure 2 for a visualization of these group results of experiments 1, 2, 3. In these analyses, only amplitude was a free parameter, since we phase-locked the fitted sinusoid such that its peak corresponded to the (removed) peaks of the phase-aligned observed data. Relevance values resulting from this curve-fitting were tested against a null distribution built from relevance values obtained by this exact same analysis performed on all 2000 trial-shuffled datasets obtained when performing the individual participant permutation tests. There was no effect of tACS phase on left hemifield contrast thresholds (R-squared = 0, P = 0.55), and a trend for right hemifield contrast thresholds (R-squared = 0.43, P = 0.08). Reaction times to left hemifield targets however were nearly significantly modulated by tACS phase (R-squared = 0.37, P = 0.0505), not right hemifield targets (R-squared = 0, P = 0.79).

**Figure 2:**
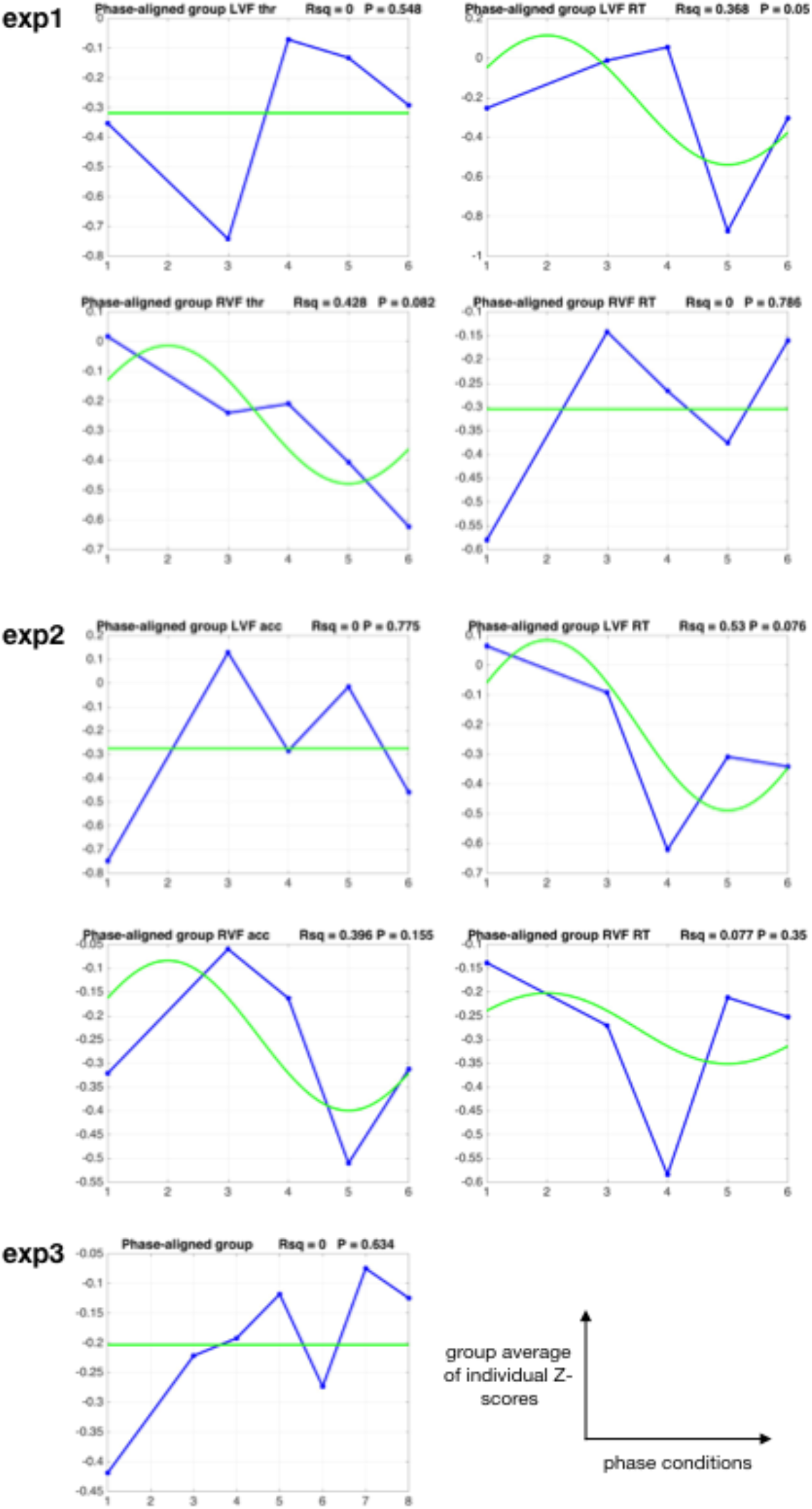
phase-aligned group average results. Individual observed results (contrast thresholds and mean RT in exp1, accuracy and mean RT in exp2, hit rate in exp3) were Z-scored, then phase-shifted such that the absolute peak value was in phase slot ‘2’ for each participant, then averaged across participants (blue lines). The group data point for phase slot ‘2’ was left out of graphs and analysis, since it was an average of the individual datapoints used for phase-alignment. tACS phase modulation of behavioral measures should result in a one-cycle sinusoidal pattern over the remaining phase conditions, with its peak at phase slot 2. Thus phase-locked best-fitting sinusoids are shown in green. Above each graph, we present the goodness of fit of these sinusoidal fits, expressed by R-squared (Rsq), and the P-value to come out of a permutation test of the associated relevance value (a measure reflecting both the variance explained and the extent of modulation, see Methods).

In a final analysis, based on this same phase-alignment procedure, a permutation test comparison of the averaged “up-phase” and “down-phase” conditions evaluated phase modulation with less of a requirement on a sinusoidal pattern (see Methods). In this analysis, tACS phase nearly significantly modulated right hemifield thresholds (P = 0.0595), not left hemifield thresholds (P = 0.98) or reaction times (left: P = 0.13, right: 0.67).

In sum, the analyses reported here paint a somewhat inconsistent picture, with essentially no support for contrast threshold modulations in left hemifield, which was the focus of this experiment. Instead, depending on the analysis, there were indications that perhaps thresholds were modulated in right hemifield. And there was evidence that reaction times to targets in left hemifield followed a sinusoidal pattern over tACS phase conditions. But given the post-hoc nature of some of these analyses, replications of these potential effects are desirable before conclusions are warranted.

### Experiment 2

#### Individual Results

Supplementary figures SF3/SF4 present individual results accuracies (SF3) and mean reaction times (SF4), with best-fitting one-cycle sinusoids superimposed. Figure titles also provide the explained variances, and the P-values based on permutation tests on the relevance values of the curve-fitting approach. Table 2 also provides the variances explained by fitted sinusoids (R-squared) for both dependent variables across conditions.

**Table 2:** individual results. Explained variance (R-squared) with P-value resulting from permutation tests on relevance values in parentheses. Hit rate analysis was the main analysis, reaction time analysis was post-hoc Bold cells are statistically significant (uncorrected)

|  | Left visual field |  |  |  | Right visual field |  |  |  |
| --- | --- | --- | --- | --- | --- | --- | --- | --- |
|  | hit rate |  | RT |  | hit rate |  | RT |  |
| 1 | **0.64** | **(0.015)** | 0.17 | (0.622) | 0.22 | (0.618) | 0.61 | (0.389) |
| 2 | 0.30 | (0.486) | **0.96** | **(0.029)** | 0.39 | (0.412) | 0.52 | (0.226) |
| 3 | 0.37 | (0.490) | 0.35 | (0.504) | 0.16 | (0.750) | 0.37 | (0.569) |
| 4 | 0.73 | (0.239) | 0.59 | (0.528) | 0.40 | (0.359) | 0.42 | (0.270) |
| 5 | 0.41 | (0.649) | 0.26 | (0.713) | 0.10 | (0.965) | 0.03 | (0.971) |
| 6 | 0.04 | (0.945) | 0.51 | (0.500) | 0.56 | (0.349) | 0.56 | (0.198) |
| 7 | 0.69 | (0.392) | 0.70 | (0.162) | 0.66 | (0.359) | 0.12 | (0.833) |
| 8 | 0.01 | (0.996) | 0.42 | (0.585) | 0.02 | (0.974) | 0.84 | (0.142) |
| 9 | 0.20 | (0.682) | 0.65 | (0.101) | 0.91 | (0.093) | 0.49 | (0.475) |
| 10 | 0.39 | (0.363) | 0.65 | (0.249) | 0.56 | (0.132) | 0.62 | (0.391) |

#### Group Results

Per hemifield, and per dependent variable, the individual P-values were converted to Z-scores, which were subsequently tested to be higher than zero in a second-level, one-sided, uncorrected t-test across participants. This test did not reject the null hypothesis that sinusoids fit the individual data consistently better than chance for accuracy (left hemifield: t(9) = −0.37, P = 0.64, right hemifield: t(9) = −0.27, P = 0.60), with barely a statistical trend for reaction time (RT) for left hemifield targets (left hemifield: t(9) = 1.45, P = 0.09, right hemifield: t(9) = 0.30, P = 0.38).

For the other group analyses, we first phase-aligned and Z-scored individual performance patterns, then removed the data points used for phase alignment, and then averaged them to create a group graph, as in exp1 (and exp3) and as shown in Figure 2. Accuracy on targets was not modulated by tACS phase (left hemifield: R-squared = 0, P = 0.78, right hemifield: R-squared = 0.40, P = 0.16). In the post-hoc analysis of reaction times, we observed a marginally significant (R-squared = 0.53, P = 0.08) effect of tACS phase on reaction times for left hemifield targets, but not right (R-squared = 0.08, P = 0.35). The permutation tests on the averaged “up-phase” and “down-phase” conditions (see Methods) yielded a significant effect of tACS phase on group average reaction times for left hemifield targets only (P = 0.047, other P’s > 0.05).

### Experiment 3

The third experiment evaluated a luminance change detection task with superior stimulus timing control, in which we recorded only hits (and misses) to calculate a hit rate per participant per phase condition (8 phase conditions).

#### Individual Results

Supplementary Figure SF5 shows the individual results, the associated best-fitting sinusoids and goodness of fits (R-squared). Relevance values based on these fits were tested against individual null distributions. For no participant in the sample of 15 did a one-cycle sinusoid explain the data significantly better than chance.

#### Group Results

For a second-level group analysis on the individual permutation results, the Z-scores corresponding to individual P-values were t-tested against zero, after removal of one statistical outlier value (inclusion did not qualitatively change the outcome). They were not significantly different from zero (t(13)=-0.33, P = 0.63), hence no evidence that tACS phase modulated luminance change detection.

Individual results (hit rates over phase conditions) were Z-scored, phase-shifted, and averaged. As shown in Figure 2, R-squared is essentially 0 and P = 0.63. Also in the final group analysis, permutation-testing the mean hit rate of the up-phase (phase conditions 1, 3, 4 in Figure 2) versus the down-phase (phase conditions 5-8) of the phase-shifted group average, resulted in no effect (P = 0.69).

## Discussion

In the current series of experiments, we implemented an advanced methodological setup (ten Oever, de Graaf, et al., 2016) to test the causal relevance of right occipital alpha-frequency tACS phase for visual processing. In experiment 1, we performed psychophysical staircases to estimate contrast thresholds for 2AFC (2-alternative forced choice) grating detection, separately and in parallel for conditions in which gratings were triggered at six pre-determined phases of the 10Hz tACS signal. In experiment 2, the tACS was administered at EEG-based individual alpha frequencies and target detection performance was tested at pseudo-fixed contrast levels. In experiment 3, a custom-made LED device was used to test potential modulation of signal detection performance (a brief LED luminance change) occurring at eight pre-determined individual alpha frequency tACS phases.

In a series of analyses, across all three experiments, no consistent or convincing tACS phase modulations of these core dependent variables could be revealed. However, depending on the analysis, there were indications that reaction times to targets in the left hemifield were modulated by tACS. These findings, obtained in both experiments that recorded reaction times, are of interest. But they are based on post-hoc analysis, and obtained in experiments that were not explicitly designed to reliably assess reaction times. This may or may not mean that the alpha tACS phase modulation of reaction times can be replicated or even more pronounced in future studies.

Alpha phase has been related to manual reaction times in EEG experiments (Callaway & Yeager, 1960; Dustman & Beck, 1965). This is in line with the ‘excitability hypothesis’ (Bishop, 1933; Lindsley, 1952), that occipital alpha phase reflects the bottom-up cortical excitability of occipital neurons. There is also evidence that frontal (high-)alpha-phase predicts saccadic reaction times (Drewes & VanRullen, 2011). For alpha power, the relation to visual efficiency has been attributed to response bias rather than sensitivity (Iemi, Chaumon, Crouzet, & Busch, 2017). In our experiment, phase effects on reaction times could reflect either of the two, though improved sensitivity might have been expected to affect the primary behavioral measures (contrast thresholds and hit rates) as well. These considerations reflect an ongoing discussion on the nature of the relation between alpha phase and visual processing. Sherman et al. (2016) for example found no relation between alpha phase and sensitivity, and rather suggested that (pre-stimulus) alpha phase reflects modulation of decision threshold by prior expectations. When it comes to positive results with tACS, one should consider recent reports that some findings could be attributable to transcutaneous/somatosensory stimulation rather than direct neural effects underneath the electrode (Asamoah, Khatoun, & McLaughlin, 2019). Future studies should evaluate to what extent tACS phase-specific effects on visual perception could be confounded by such indirect effects as well as peripheral phosphenes.

For our dependent variables of a priori interest; contrast thresholds, hit rates, and signal detection rates, our null results can be explained in several ways. Firstly, perhaps tACS did not successfully phase-align neuronal alpha oscillations, and while naturally occurring alpha phase is functionally relevant, alpha tACS phase is not. While alpha tACS effects on EEG alpha power have been reported repeatedly (Kasten, Dowsett, & Herrmann, 2016; Neuling, Rach, & Herrmann, 2013), it is possible that mechanisms other than phase-alignment (i.e. stochastic resonance, Vossen, Gross, & Thut, 2015) underlie some of those effects. Secondly, perhaps the phase of natural alpha oscillations occurring at lower right occipital cortex is not functionally relevant. Using TMS, Jaegle and Ro (2014) could show that the phase of an alpha train was functionally relevant when the TMS was applied over parietal cortex, but not when applied over occipital cortex. Thirdly, perhaps the problem is in the dependent variables. In the context of alpha power, Lange et al. (2013) showed that alpha power indexes enhanced cortical excitability, not improved visual perception. One might ask the same question about alpha phase, as discussed above. Contrast thresholds and hit rates might better capture visual acuity while reaction times better capture excitability. Lastly, it is of course possible that our null results are trivial; perhaps some aspects of our methodology were suboptimal to revealing tACS phase modulations which could have been obtained with larger samples or different experimental setup or design. We did validate the experimental approach itself (ten Oever, de Graaf et al., 2016), and recently used it to demonstrate beta-frequency tACS phase effects on motor-evoked potentials (Schilberg et al. 2018).

## Conclusion

It remains difficult to interpret null results in NIBS (see de Graaf TA & Sack, 2018; de Graaf & Sack, 2011). The collection of null results presented here are inconclusive in our view (Level C-B null evidence, see de Graaf & Sack, 2018). Yet they seem worthy of dissemination to guide future studies, share the advanced experimental procedures and analysis approaches, and for the sake of transparency in a growing literature of alpha power and phase studies. The post-hoc positive results for tACS phase modulation of left hemifield reaction times are promising, and not unexpected, but should be interpreted with caution for reasons outlined above.

## Data/Code availability

Data and code will be made available upon reasonable request.

---

**SF1:**
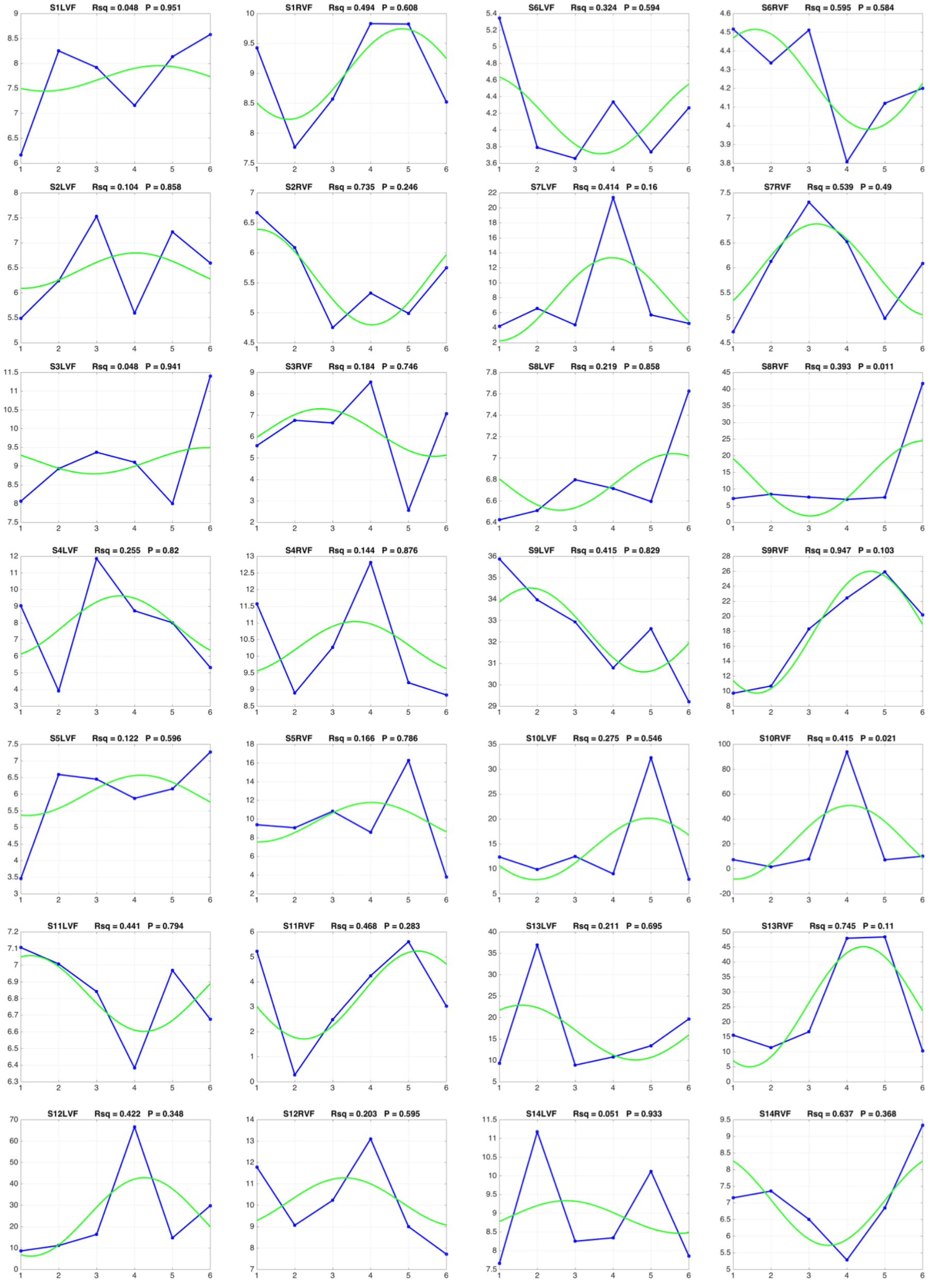
individual results experiment 1: thresholds

**SF2:**
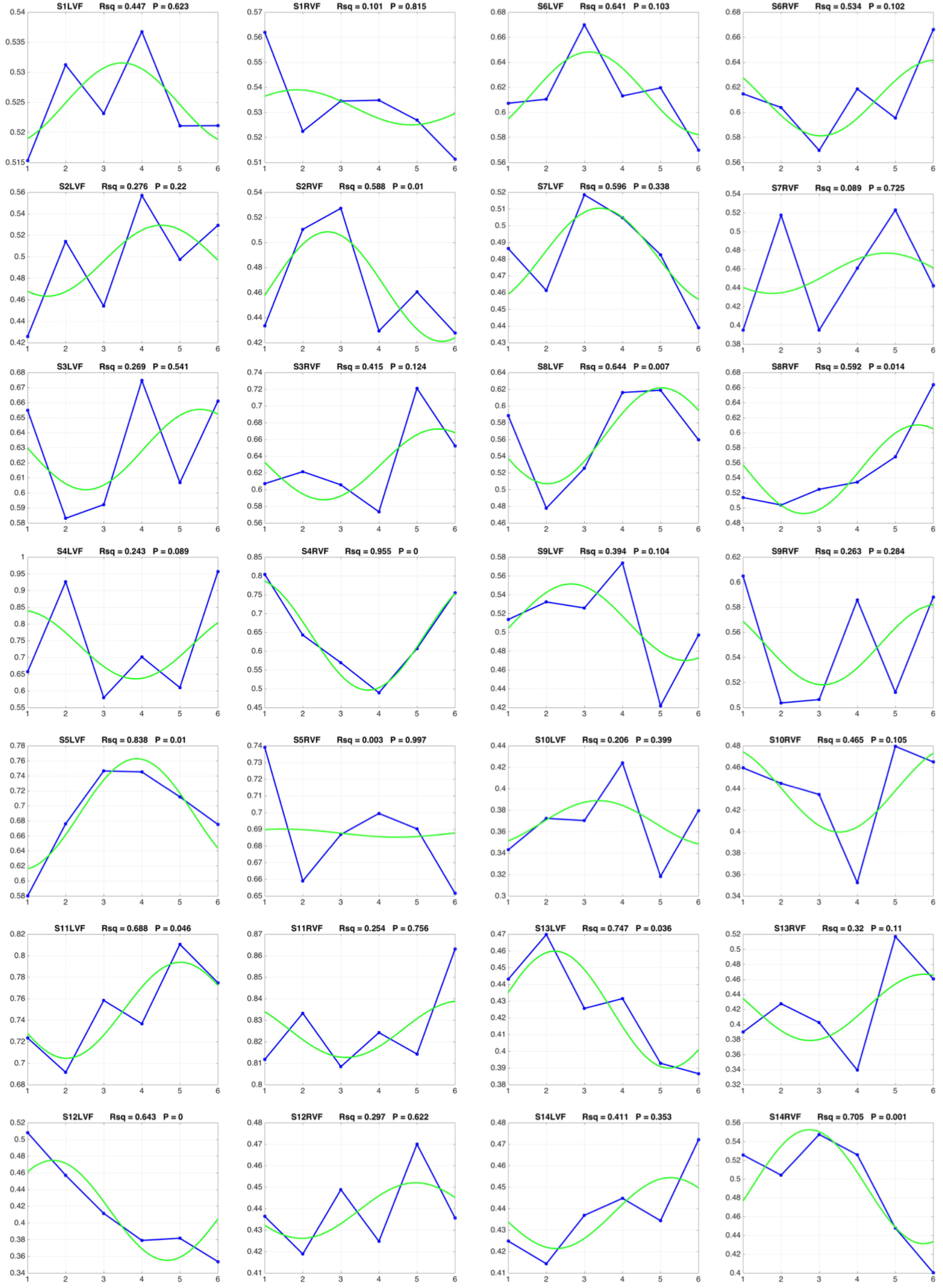
individual results experiment 1: reaction times

**SF3:**
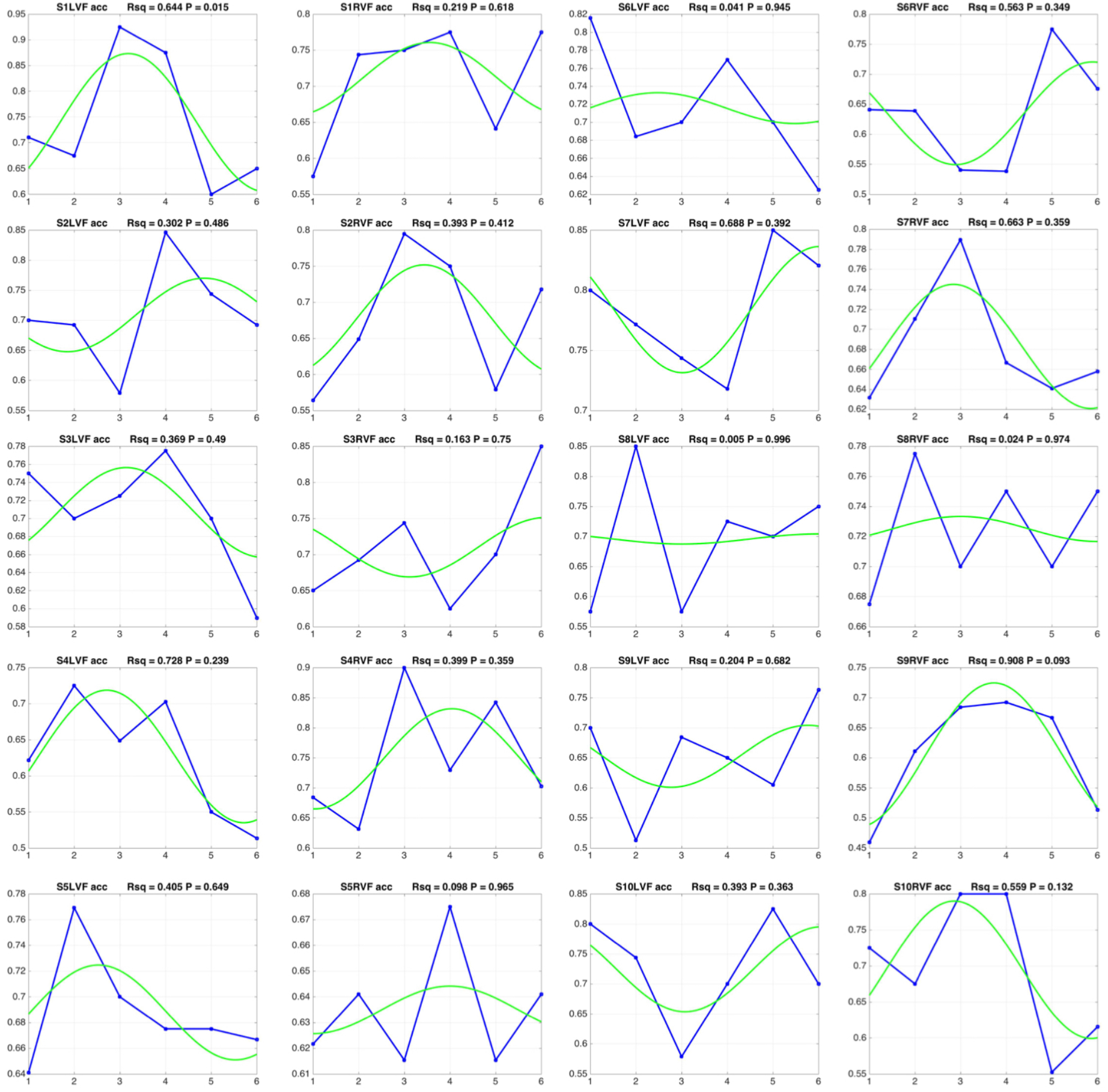
individual results experiment 2: accuracy

**SF4:**
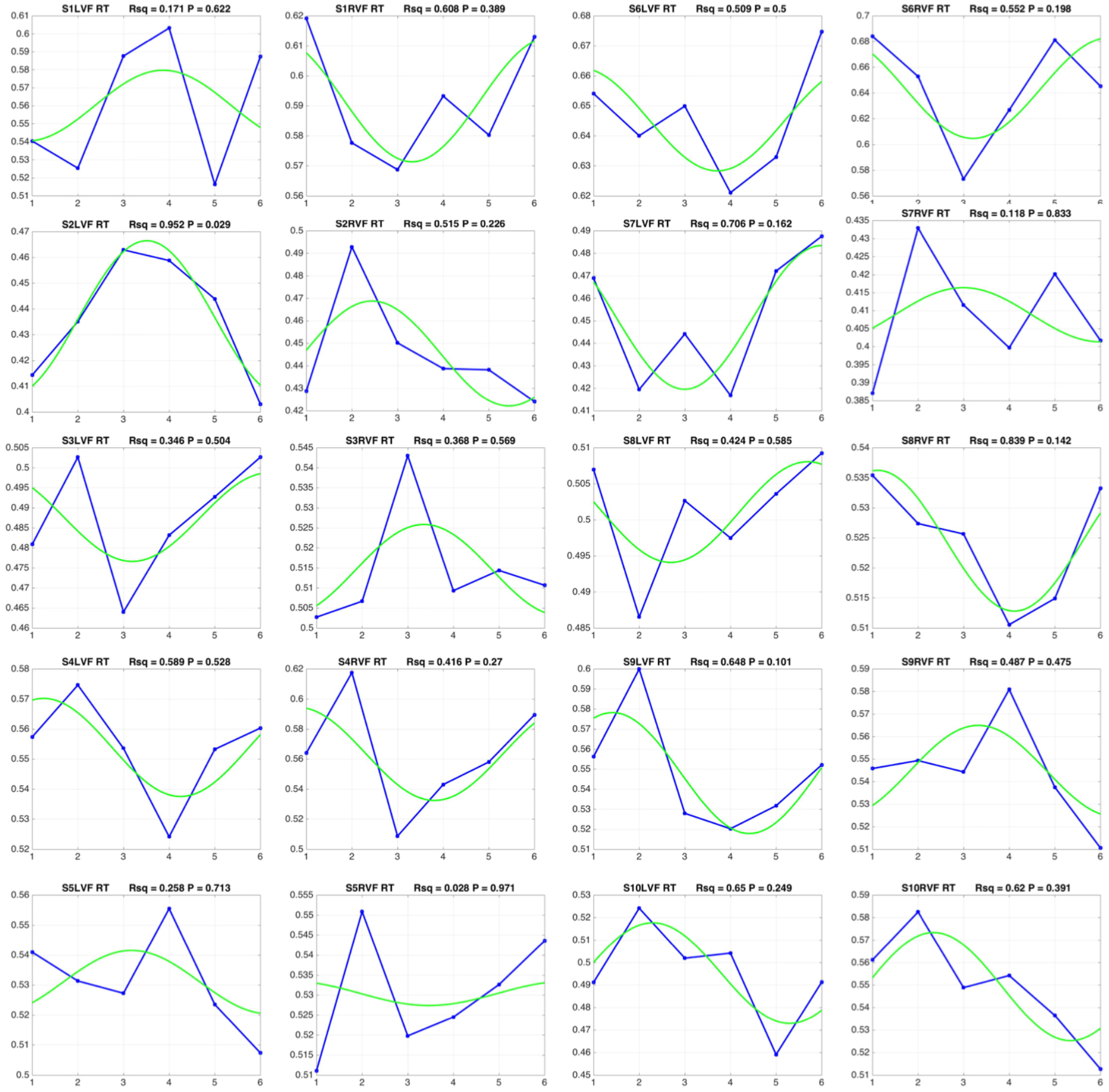
individual results experiment 2: reaction times

**SF5:**
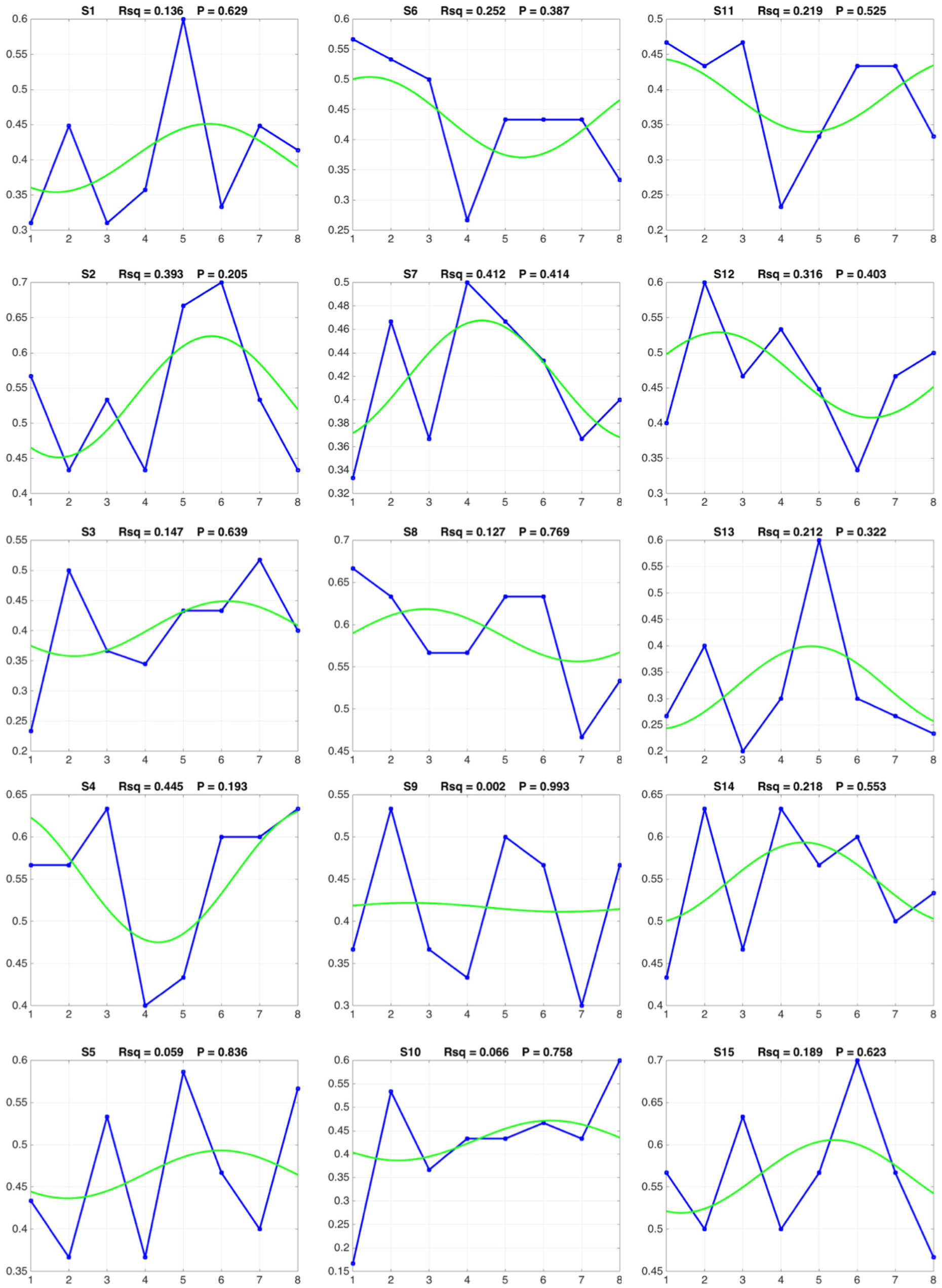
individual results experiment 3: detection rates

